# A synergistic generative-ranking framework for tailored design of therapeutic single-domain antibodies

**DOI:** 10.1101/2025.05.09.653014

**Authors:** Yu Kong, Jiale Shi, Fandi Wu, Ting Zhao, Rubo Wang, Xiaoyi Zhu, Qingyuan Xu, Yidong Song, Quanxiao Li, Yulu Wang, Xingyu Gao, Yuedong Yang, Yanling Wu, Zhenlin Yang, Jianhua Yao, Tianlei Ying

## Abstract

Single-domain antibodies (sdAbs) have emerged as powerful therapeutic agents due to their small size, high stability, and superior tissue penetration. However, unlike conventional monoclonal antibodies (mAbs), sdAbs lack an Fc domain, limiting their functional versatility and manufacturability. To address this challenge, we developed TFDesign-sdAb, a deep learning-based generative-ranking framework that enables rational engineering of sdAbs with tailored functionalities. Our framework integrates a structure-aware diffusion model (IgGM) for large-scale candidate generation and a fine-tuned sorter (A2binder) that evaluates and prioritizes them based on predicted functionality. Unlike traditional CDR-focused approaches, TFDesign-sdAb optimizes both complementarity-determining regions (CDRs) and framework regions (FRs), allowing sdAbs to acquire new functional properties while maintaining antigen specificity. We validated our approach by conferring Protein A binding to human VHs and nanobodies that originally lacked this feature, achieving high expression rates, strong binding affinities, and successful purification via industry-standard Protein A affinity chromatography. High-resolution structural characterization (1.49 Å and 2.0 Å) of the redesigned sdAb-Protein A complexes revealed conserved FR-mediated binding motifs that recapitulate natural Fc-Protein A interactions, validating the accuracy of our model. Furthermore, our pipeline streamlined the antibody engineering process, achieving a 100% success rate in generating Protein A-binding sdAbs while maintaining their original antigen-binding affinity. This work demonstrates the power of AI-driven design in overcoming long-standing limitations in antibody engineering and presents a scalable, generalizable solution for enhancing sdAb functionality.

## Introduction

Single-domain antibodies (sdAbs), including camelid-derived nanobodies and human-derived variable heavy domains (VH), have emerged as a transformative class of biologics due to their small size, high stability, and superior tissue penetration^1^. Compared to conventional monoclonal antibodies (mAbs), sdAbs offer superior accessibility to sterically restricted epitopes, enable cost-effective, high-yield production in microbial systems, and are well-suited for non-invasive delivery methods, such as aerosolized administration^2^. However, the lack of an Fc domain in sdAbs results in the loss of certain functionalities mediated by Fc-dependent protein interactions^3–5^. For instance, a critical challenge for the therapeutic application of sdAbs is their intrinsic lack of Protein A binding, which prevents the use of the industry-standard, tag-free purification process established for mAbs^6^. To enhance their manufacturability and therapeutic potential, tailored design strategies are needed to endow sdAbs with specific functionalities while preserving their antigen affinity.

Engineering sdAbs with new functionalities remains a labor-intensive and inefficient process. Traditional methods rely on combinatorial library screening, such as phage display, to identify variants with desirable properties^7,8^. These approaches require extensive rounds of biopanning and typically yield low success rates, making them impractical for large-scale or systematic functionalization^9^. Furthermore, each sdAb must be independently optimized through this resource-intensive process, with most attempts resulting in failure. Therefore, although a few sdAbs have been reported to exhibit weak Protein A affinity through their framework regions (FRs), these isolated instances have not resulted in a generalizable strategy for rationally engineering Protein A binding into sdAbs^10,11^.

Deep learning has revolutionized protein engineering, with generative models achieving remarkable success in antibody design by co-optimizing sequences and structures^12,13^. Recent advances, such as diffusion-based models and protein language models, have facilitated complementarity-determining regions (CDR) optimization for antigen binding^14^. However, these methods are largely constrained to CDR-focused design and do not account for FR-mediated interactions, which are essential for engineering new functionalities like Protein A binding^15^. Moreover, most deep learning frameworks rely heavily on large datasets, making them impractical for cases where experimental data are scarce^16^.

To address this challenge, we developed TFDesign-sdAb, a synergistic generative-ranking framework integrating a generative protein design model (IgGM) with a structure-informed ranking model (A2binder). By leveraging high-resolution structural insights and a small set of experimentally validated affinity data, we established a data-efficient paradigm for tailoring sdAbs with novel binding properties while preserving antigen specificity. Using this approach, we successfully reprogrammed multiple human-derived VHs and camelid-derived nanobodies to acquire high Protein A affinity, enabling efficient, tag-free purification via industry-standard Protein A affinity chromatography. High-resolution structures (1.49 Å and 2.0 Å) of the redesigned nanobodies complexed with Protein A revealed that their interaction pattern closely resembles that of the VH-Protein A complex used for model training, validating our method and demonstrating its potential to guide the rational design of sdAbs with tailored binding properties. This strategy offers a generalizable approach to functionalizing sdAbs, accelerating their development as next-generation biologics.

## Results

### Designing sdAbs with TFDesign-sdAb

Traditional antibody engineering methods, such as phage display or yeast display, involve introducing mutations into the initial antibody to generate variant libraries. These approaches require multiple rounds of selection, amplification, and screening to identify candidates with the desired properties (Fig. 1a). However, this process is labor-intensive and time-consuming, and more critically, the introduced mutations often disrupt the antibody’s original affinity for its target antigen. This makes it challenging to preserve specificity while introducing new functionalities, particularly in the context of tailored sdAb design, which requires precise modifications.

**Fig. 1.**
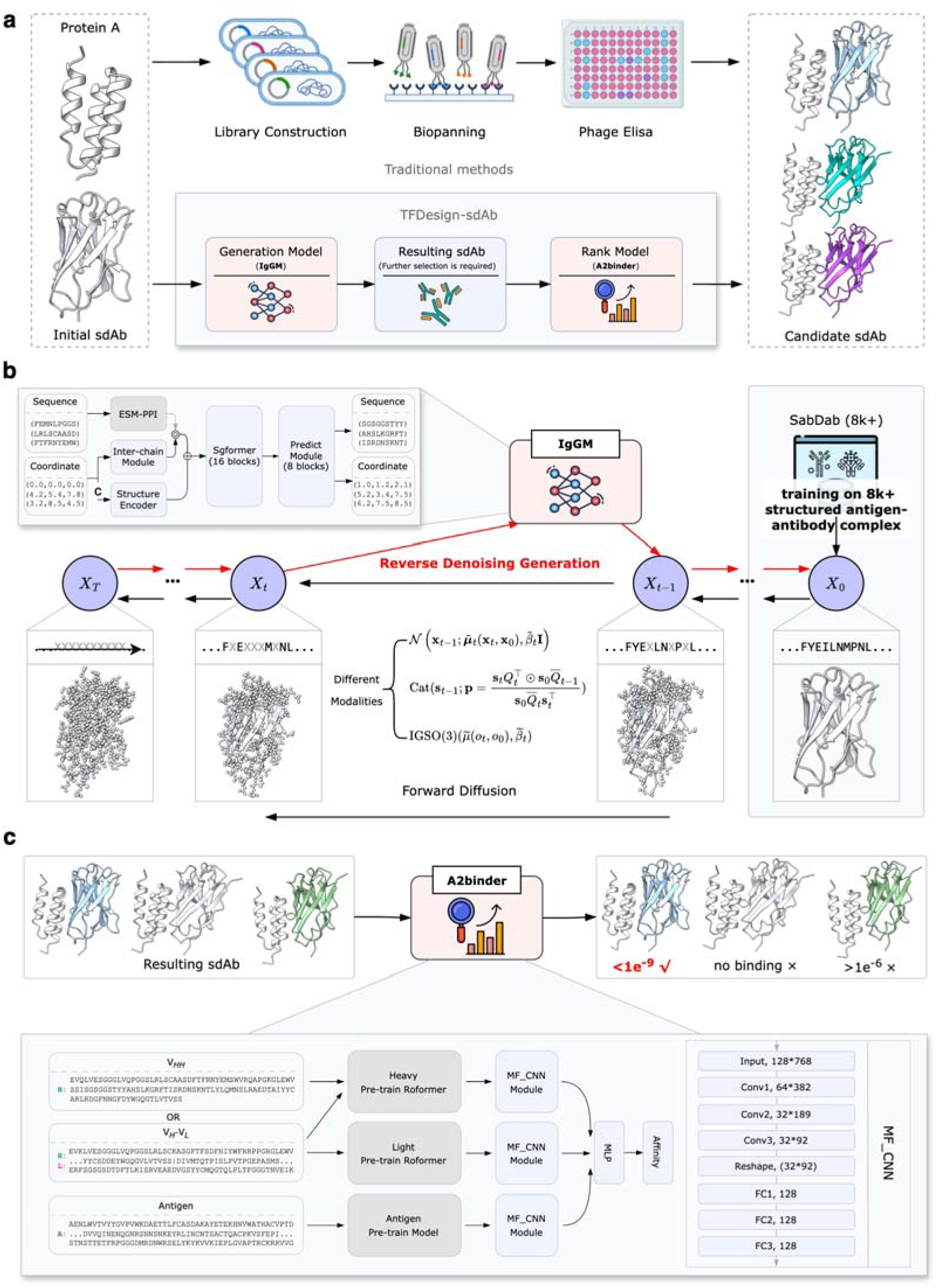
Comprehensive pipeline for tailored sdAb design. **a.** Traditional sdAb engineering methods involve complex processes, such as combinatorial library construction, screening, and identification. TFDesign-sdAb generates a batch of sdAbs through an integrated generative model, followed by ranking and screening via a scoring model to obtain candidate antibodies. **b.** The generative model of TFDesign-sdAb is based on IgGM, a hybrid-modality diffusion generative model that iteratively generates accurate sdAb sequences and structures through multiple refinement steps. **c.** The ranking model of TFDesign-sdAb is based on A2binder, which integrates pre-trained antigen/antibody large language models and employs a convolutional neural network to predict binding affinity, thereby enabling the ranking of candidate sdAbs.

To overcome these limitations, we propose a deep learning-based generative-ranking framework capable of directly predicting and designing sdAbs with targeted functionalities, reducing the need for extensive mutation cycles and ensuring the preservation of antigen specificity. As shown in Fig. 1a, our deep learning-based framework, TFDesign-sdAb, consists of two main components: a generator called IgGM and a ranker called A2binder. The IgGM model utilizes the known structure of the functional target (e.g., Protein A), along with the epitope and the sdAb sequence requiring modification, to generate a substantial number of candidate sdAbs. The A2binder model subsequently scores and ranks all candidate sdAbs, with the top-ranked candidates selected for wet lab validation.

First, we developed IgGM as a purpose-built diffusion model specifically for single-domain antibody (sdAb) functional engineering. ^17^ This generative tool enables co-design of sdAb sequences and structures based solely on partial sequences and the epitope of the antigen, without necessitating additional information such as the structure of the antibody-antigen complex. Fig. 1b depicts the forward and reverse processes involved in the co-design of antibody sequences and structures using the diffusion model component of IgGM. The network structure of IgGM consists of three parts. One part is in the form of initializing coordinates and sequences as features. Specifically, it contains a pre-trained language model ESM-PPI for extracting evolutionary features of multiple chains. The three-dimensional coordinate features are initialized through the Structure Encoder and Inter-chain module. After that, these features will be input into SgFormer for feature fusion. Finally, the fused features will be input into the Predict Module to predict the updated sequences and structures. During the training phase, noise is introduced to the structures and sequences of antibodies, allowing the model to predict and recover accurate structures and sequences. In the sampling phase, the model begins with initial noisy sequences and structures, predicts the modified sequences, and generates complete structures. This iterative process facilitates the refinement and enhancement of antibody designs. Due to the inherent properties of consistency models^18^, IgGM supports single-step generation, enabling the rapid and large-scale production of candidate antibody sequences and structures. Furthermore, it accommodates various edits and applications in zero-shot editing without the need for explicit training, thereby seamlessly integrating with other tools to enhance the quality of the generated outputs.

Next, we further enhanced the IgGM framework with a topology-aware training strategy to enable simultaneous optimization of both FRs and CDRs. Most existing antibody design methods primarily focus on the design of CDRs and do not accommodate modifications in FRs. The original IgGM only supports the design of the antibody CDR region. To achieve FR-mediated functionalization without compromising antigen affinity, we engineered IgGM with a two-phase training architecture specifically tailored for sdAb framework optimization. In the first phase, we concentrated on structural prediction tasks utilizing all antibody-antigen complex pairs from SAbDab^19^, without introducing noise to the sequences. The second phase integrated both sequence and structural prediction, focusing specifically on the antibody FR regions that interact with the antigen. This targeted approach allowed for effective optimization of the sdAb FR regions. Further details can be found in the Methods section.

To demonstrate the application of our model in functionalizing sdAbs, we focused on imparting Protein A-binding capability to sdAbs that originally lack this functionality. Protein A-based affinity chromatography is widely regarded as the gold standard for mAb purification, ensuring high efficiency and purity^20^. Derived from *Staphylococcus aureus*, Protein A is a 42 kDa surface protein consisting of five homologous immunoglobulin-binding domains (A-E) and an S region that functions as a signal sequence and cell wall-binding region (XM)^20,21^. It strongly interacts with the CH2-CH3 region within the Fc domain of IgG1, IgG2, and IgG4 and can also bind certain antibody fragments^21–24^. Although most sdAbs do not naturally bind Protein A, we had previously identified an sdAb with this capability which provided a valuable opportunity to incorporate structural insights into the model and test its predictions^2^.

During the sampling process of IgGM, we input the sdAb to be modified along with the structure of Protein A (and its epitope), resulting in the generation of a large pool of candidate sdAbs. Following the generation of candidate sdAbs, we employed A2binder to predict their affinity for Protein A. A2binder is an affinity prediction model based on a pre-trained protein language model (PLM), and we ranked the candidates according to their predicted affinities. Fig. 1c shows the model architecture of A2binder. The input of A2binder is a pre-trained antigen language model, an antibody heavy chain language model and an antibody light chain language model. It extracts the learned evolutionary knowledge. This knowledge can help subsequent modules better learn the interactions between different antigens and antibodies. After that, these features will undergo feature fusion through MF_CNN and MLP to learn interaction and output affinity values. The network architecture of MF_CNN is shown at the bottom right of Fig. 1c. It contains multiple layers of convolutional neural networks and fully connected layers for feature aggregation. It should be emphasized that the original A2binder model was not specifically engineered to handle sdAb data.

To overcome this limitation, we fine-tuned the pre-trained A2binder model by utilizing all affinity data collected from SAbDab, in conjunction with a small set of experimentally measured Protein A affinity data for our sdAb and its variants, as elaborated in the Methods section. This adaptation enables A2binder to effectively predict the affinities of sdAbs, allowing us to select the most promising candidates based on their predicted binding strength to Protein A.

### IgGM could accurately model the sdAb-Protein A interaction

To assess the reliability of the re-architected IgGM generative model, we first used it to predict the structures of the sdAb and its complex with Protein A, then experimentally determined the high-resolution X-ray crystallography structures of the same complex and conducted a comparative analysis of the predictions and experimental data. The sdAb n501 is a fully human single-domain antibody we previously identified that targets the oncofetal antigen 5T4^25^. It exhibits nanomolar affinity for Protein A and can be effectively purified using commercial Protein A resins, achieving higher purity and comparable yields to IMAC in a single step (Fig. S1). To elucidate the binding mode of n501 to Protein A, we first employed IgGM to predict the structure of the n501-Protein A complex. As illustrated in Fig. 2a, we utilized the known mAb-Protein A complex structure (PDB_ID: 1dee)^26^ to determine the structure of Protein A and its epitope when bound to the mAb. Subsequently, we input the sequence of n501 and Protein A epitope into IgGM to predict the n501-Protein A complex structure. From the IgGM-predicted complex, we identified that the paratope of n501 is located within the FR1 and FR3.

**Fig. 2.**
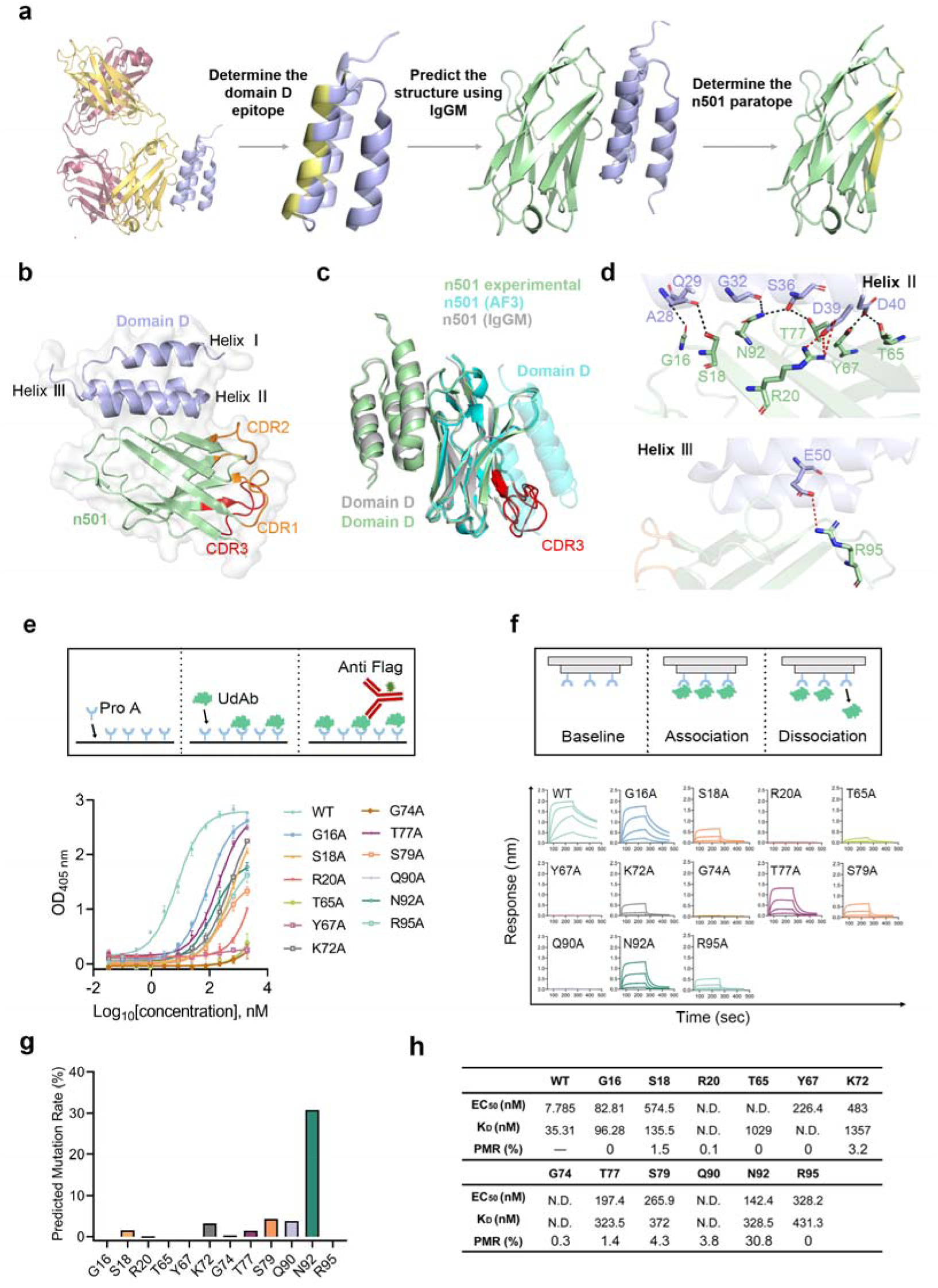
Accurate prediction of the sdAb-Protein A interaction using IgGM. **a.** Illustration of the complex structure between the n501 and domain D of Protein A, as predicted by IgGM. The domain D is depicted in purple, binding epitope of domain D in yellow, and sdAb n501 in green. **b.** The crystal structure of sdAb n501 in complex with domain D of Protein A (3.57 Å), with n501 colored in pale green and domain D in purple. The three CDRs are highlighted: CDR1 and CDR2 in orange, and CDR3 in red. **c.** Comparison of the n501-domain D structure derived from experimental data, AlphaFold 3 predictions, and IgGM predictions is presented. The experimentally determined structure is colored in pale green, AlphaFold 3 predictions in cyan, and IgGM predictions in grey. CDR3 of n501 is marked in red. **d.** The polar interactions between n501 and domain D are shown, with all interacting residues shown as sticks, colored pale green for n501 and light blue for domain D. Hydrogen bonds are indicated with black dashes, and salt bridges are shown in red. **e.** Binding activity of n501 variants with domain D, as measured by ELISA. A schematic diagram of the assay is shown in the upper panel, and the concentration-response curve is presented in the lower panel. Three replicates were performed for each experiment. **f.** Binding affinity of n501 variants with domain D, as measured by BLI. A schematic diagram of the experimental procedure is shown in the upper panel, with the binding kinetics curves presented in the lower panel. **g.** Predicted mutation rate of sequence generated by IgGM across 12 positions, ased on an analysis of 1000 variants. **h.** Summary table for ELISA binding assay, BLI binding assay and predicted mutation rates.

Next, we co-crystallized n501 with the domain D of Protein A and successfully determined their complex structure at a resolution of 3.57 Å (Fig. 2b). We then compared the experimentally determined n501-Protein A complex structure with those generated by IgGM and AlphaFold-3. As shown in Fig. 2c, IgGM accurately predicts the interaction between n501 and Protein A (DockQ=0.834, RMSD=1.29 Å), whereas AlphaFold-3 incorrectly docked Protein A to the complementarity determining region (CDR) (DockQ=0.028, RMSD=13.11 Å). This demonstrates that our fine-tuned IgGM model significantly outperforms in accuracy and reliability when predicting the interactions of sdAbs with Protein A.

Analysis of the crystal structure reveals that the interaction between n501 and domain D of Protein A is predominantly mediated by FR1 and FR3. This was further supported by the observation that two chimeric sdAbs where the CDRs are grafted onto the framework regions of n501 maintained robust binding to Protein A (Fig. S2). Specifically, FR1 and FR3 were found to interact with helices II and III of Protein A domain D (Fig. 2d). Further analysis indicated that 12 residues within n501 are involved in this interaction, including G16, S18, R20 from FR1, and T65, Y67, K72, G74, T77, S79, Q90, N92, R95 from FR3 (Supplementary Table 1, IMGT numbering). Notably, six of these residues are located on the β-strand, while the others are situated on the interstrand loops opposite the CDRs. The domain D of Protein A contributes an additional 12 residues to the binding interface, including A28, Q29, G32, F33, Q35, S36, D39 from helix II, N46, V47, E50 from helix III, and K53 from the interloop connecting the two helices. Seven hydrogen bonds are formed between n501 and Protein A (G16-Q29, S18-A28, R20-D39, T65-D40, Y67-D40, T77-S36, N92-G32/S36), along with two salt bridges (R20-D39 and R95-E50). Additionally, residues K72, G74, S79, and Q90 of n501 primarily interact with Protein A through hydrophobic interactions.

To assess the role of these amino acids, we conducted an alanine scan and systematically evaluated the impact of these 12 residues using two methods (Fig. 2e). ELISA results indicated that all 12 alanine-substituted mutants exhibited decreased affinities for Protein A, suggesting a collective contribution of these residues to the interaction (Fig. 2f). Notably, mutations R20A, T65A, Y67A, G74A, and Q90A displayed the most significant negative effects, a trend further validated by biolayer interferometry (BLI).

We also verified the accuracy of the key amino acids predicted by IgGM. Using IgGM, we generated 1000 variant sequences and analyzed the mutation probabilities at these 12 positions, as shown in Fig. 2g. The mutation probabilities for R20, T62, Y67, and G74 were found to be 0.1%, 0.0%, 0.0%, and 0.3%, respectively, underscoring the critical importance of these positions for the binding of n501 to Protein A. This supports the accuracy of IgGM in predicting not only the n501-Protein A complex structure but also in identifying most of the key binding residues from both structural and sequence perspectives. However, some discrepancies were noted; for instance, Q90 exhibited a 3.8% probability of mutating to other amino acids. While predicted mutation probabilities can indicate key binding residues, they do not always correlate with changes in binding affinity (ΔΔG). For example, N92 had a predicted mutation probability as high as 30%, yet the mutation of asparagine to alanine at this position resulted in a binding affinity decrease from 35.3 nM to 328.5 nM, a more significant change than that observed for mutations such as G16A and S18A. Therefore, the development of an affinity ranker is essential. The binding affinities before and after mutations, along with the predicted mutation probabilities, are summarized in Fig. 2h.

### Endowing sdAbs with Protein A binding capacity

To evaluate the capability of TFDesign-sdAb in endowing previously non-binding sdAbs with the ability to bind Protein A, we initially selected three fully human VHs for validation: n622, n67, and n118. These antibodies target distinct antigens—CEACAM5, Zika E protein, and human CD16A, respectively. For each initial sdAb, we utilized IgGM to predict the structure of the sdAb-Protein A complex and determined the paratope of the sdAb based on this structural information, as illustrated in Fig. 2a. We then employed IgGM to modify these positions, generating 1000 candidate variants for each sdAb while excluding those with a Levenshtein distance greater than 3, as excessive modifications could adversely affect the binding affinity between the sdAbs and their respective antigens.

Subsequently, we ranked the remaining candidates and selected the top 5 based on their predicted affinity scores. The affinity of these selected sdAbs for Protein A was quantified using BLI. The results indicated that all mutants were successfully expressed (Fig. S3), with four out of five mutants from n622 and n67, and three from n118, demonstrating binding to Protein A, yielding an overall success rate exceeding 60%. The affinities of these mutated sdAbs for Protein A ranged from tens to hundreds of nanomolar, thereby validating the reliability of TFDesign-sdAb (Fig. 3a-c).

**Fig. 3.**
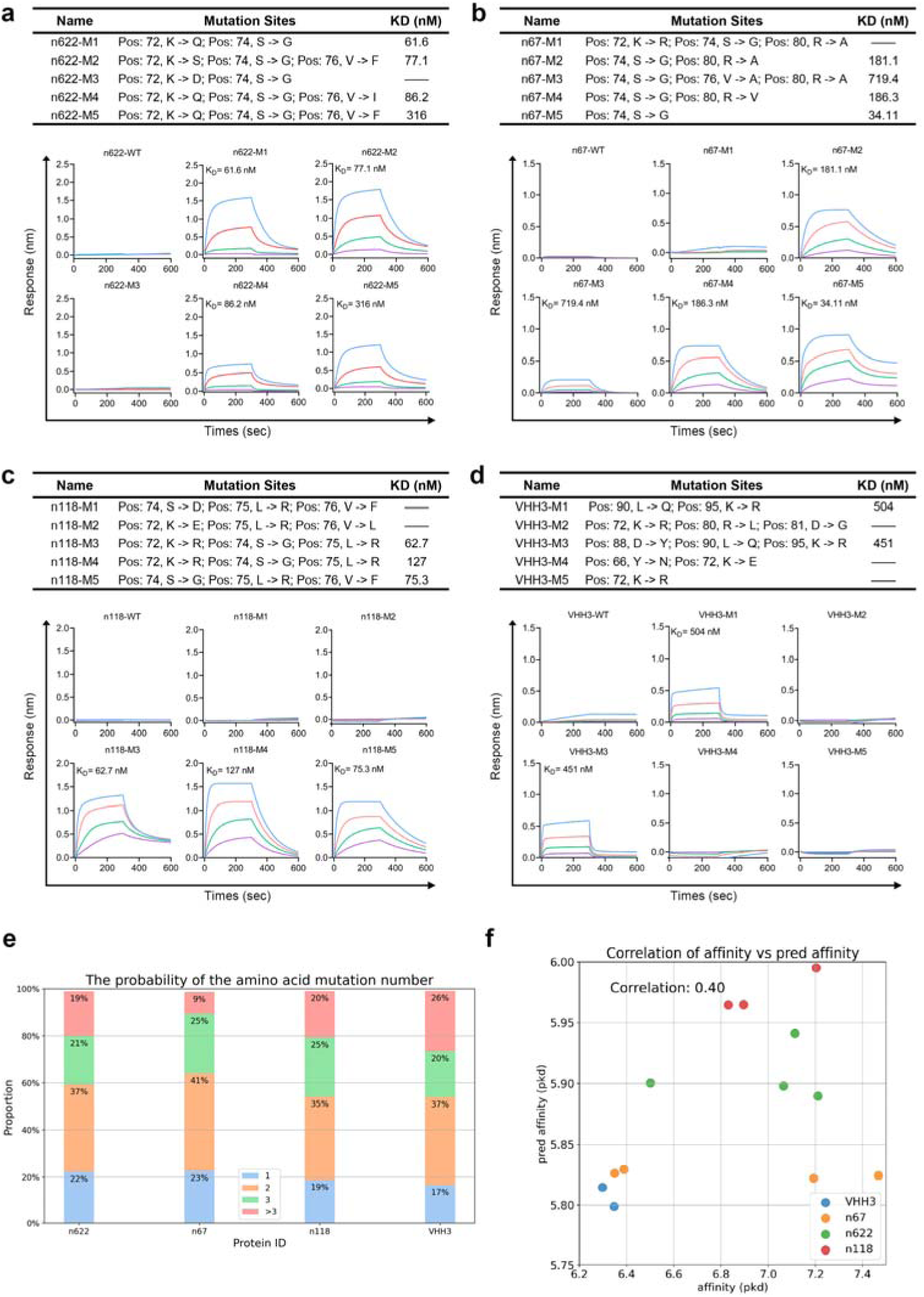
Endowing Protein A binding capability to sdAbs. **a-d.** Four sdAbs, including three human-derived (n622, n67, and n118) and one camel-derived VHH (VHH3), which lacked Protein A binding activity, were selected. For each sdAb, 1000 candidate variants were generated using the model. The top 5 variants, based on predicted Protein A-binding affinity, were selected and quantified by BLI. **e.** Proportion of mutations among 1000 sequences generated by the IgGM model, color-coded by mutation count. **f.** Correlation analysis between predicted and experimentally measured KD values for sdAb candidate variants.

Additionally, we selected the camelid nanobody VHH3, which targets human TNFα, shares low sequence similarity with n501, and does not bind to Protein A, to further evaluate the versatility of our model. For VHH3, all five generated VHH candidates were successfully expressed, with two demonstrating the ability to bind Protein A (Fig. 3d), exhibiting affinities of 504 nM and 451 nM, respectively.

We also analyzed the candidates generated by IgGM. The number of mutations in the sequences produced by IgGM predominantly ranged from 1 to 2, as illustrated in Fig. 3e. This observation suggests that the FRs of the sdAbs are relatively conserved. Additionally, we compared the predicted affinity values generated by A2Binder with actual affinity measurements using 13 binding sdAbs that had quantifiable affinity results. The dissociation constant (*Kd*) values were transformed

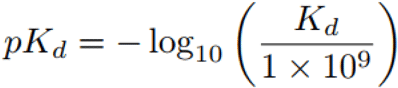

into log space, denoted as *pKd*, following the methodology outlined in He et al. [2017], as detailed in Eq. 1 to facilitate data smoothing.

As shown in Fig. 3f, the Pearson correlation coefficient between the predicted affinity values and the actual affinity values was found to be 0.4, indicating a moderate degree of correlation; however, this correlation was not statistically significant. Notably, when A2Binder was used without incorporating sdAb affinity data for fine-tuning, the Pearson correlation on the same test set was −0.36. This result highlights A2Binder’s limitations in addressing the zero-shot problem. In contrast, after fine-tuning with a small amount of sdAb data, A2Binder’s performance exhibited a significant improvement.

### Structural characteristics of redesigned sdAbs

To investigate the mechanism by which a limited number of mutations can confer Protein A binding to sdAbs, we analyzed the effects of various mutations from a structural perspective. Our analysis revealed that in the three sdAb mutants, the mutation sites were predominantly located in the region spanning amino acids 72 to 76 (Fig. 4a). This region is situated on the loop of FR3 that connects β-strands C’’ and D. Notably, all successfully optimized mutants contained the S74G mutation. In the case of n501, the glycine at position 74 primarily forms hydrophobic interactions with Protein A. However, mutating this site to serine or aspartic acid could introduce steric hindrance with helix III of Protein A domain D. Furthermore, we observed that mutating this site to alanine, which has a methyl side chain, also significantly impacts binding to Protein A (Fig. 2e-f). Only glycine, possessing the smallest side chain, can effectively prevent steric hindrance, indicating stringent amino acid constraints at this position. Additionally, by comparing five mutants of n67, we found that the amino acids surrounding the 74th position also influence the binding affinity of Protein A, suggesting that the region encompassing amino acids 72 to 76 is critical for the interaction between sdAbs and Protein A.

**Fig. 4.**
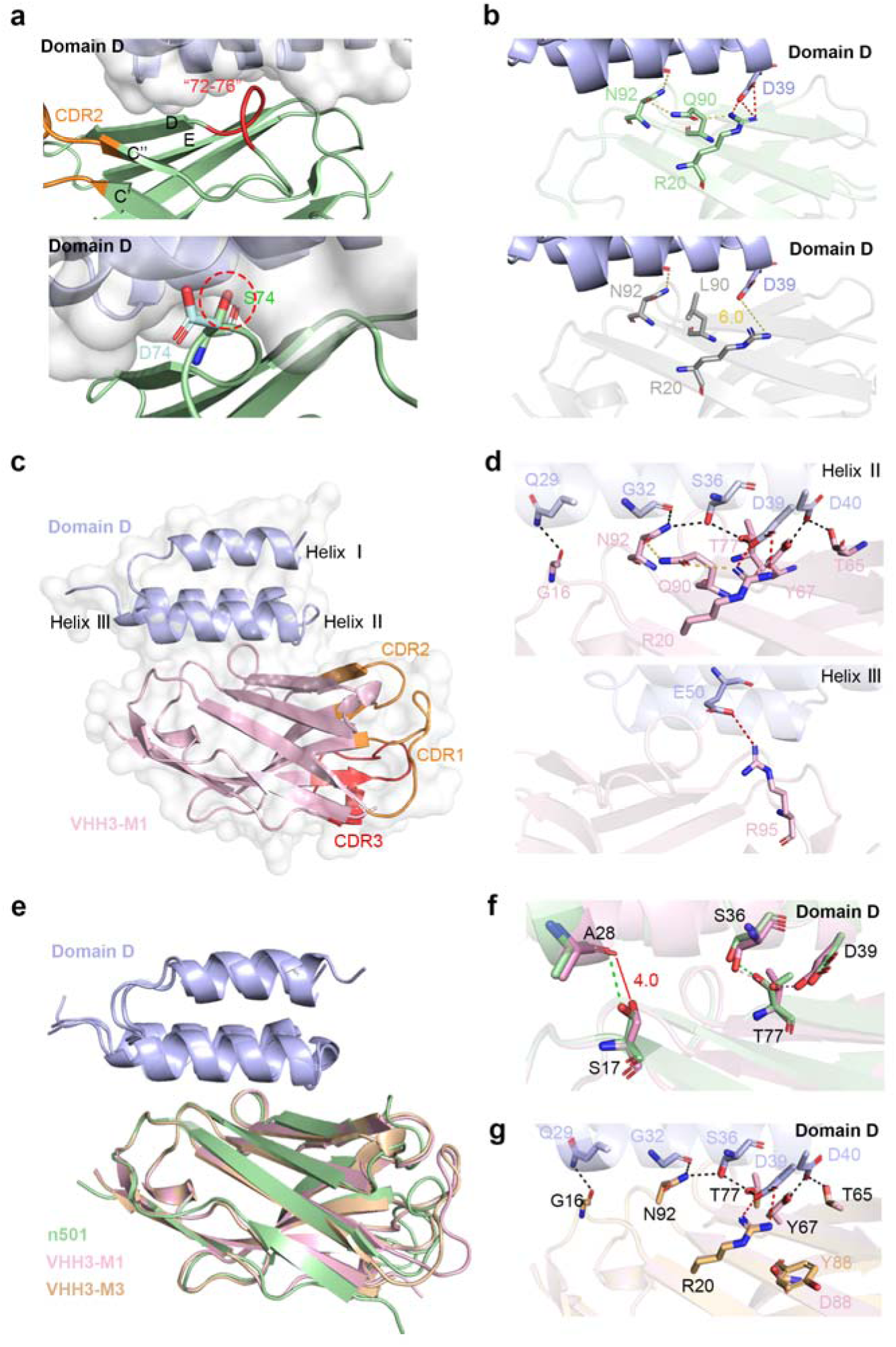
Structural characteristics of optimized sdAb. **a.** Structural analysis of key binding sites between human-derived sdAb and Protein A. The 72-76 amino acid residues are highlighted in red, with surrounding β-strands numbered according to previous reports (upper panel). The 74th site mutation to serine (green) or asparagine (cyan) causes steric hindrance with domain D, marked by red circles (lower panel). Amino acid numbering is based on the IMGT nomenclature. **b.** Structural analysis of key binding sites between camel-derived VHH and Protein A. In the upper panel, the interaction between glutamine at position 90 and residues N92 and R20, as well as the interaction between R20 and domain D, is depicted. In the lower panel, the potential interactions between leucine at position 90 and residues N92 and R20, as well as the potential interaction between R20 and domain D, are illustrated. Hydrogen bonds are represented by yellow dashed lines, and salt bridges are indicated by red dashed lines. **c.** The crystal structure of VHH3-M1 and domain D of Protein A (1.49 Å), with VHH3-M1 represented in pink and domain D in light blue. The three CDRs are marked in orange or red, respectively. **d.** The polar interactions between VHH3-M1 and domain D. All interactive residues are shown in stick and colored in pink for VHH3-M1 and light blue for domain D. The hydrogen bonds are shown in black dashes while salt bridge in red. **e.** The comparison of the complex structures between n501, VHH3-M1, and VHH3-M3 with domain D is depicted. Domain D is represented in light blue. n501, VHH3-M1, and VHH3-M3 are displayed in pale green, pink, and orange, respectively. **f.** The comparison of the binding of VHH3-M1 and n501 to domain D is illustrated. The interacting amino acids are displayed as sticks. Pink dashed lines represent the hydrogen bond interactions between VHH3-M1 and domain D, while green dashed lines indicate the hydrogen bond interactions between n501 and domain D. **g.** The comparison of the binding of VHH3-M1 and VHH3-M3 to domain D is presented. The interacting amino acids are depicted as sticks, with residues in VHH3-M1 represented in pink and those in VHH3-M3 in orange. Domain D is depicted in light blue.

We also identified the L90Q mutation in both VHH3-M1 and VHH3-M3. The glutamine at position 90, similar to glycine at position 74 in n501, forms hydrophobic interactions with Protein A. However, its side chain can form hydrogen bonds with the side chains of N92 and R20, which are essential for binding to Protein A (Fig. 2e-f). This interaction can draw the side chain of R20 closer to D39 of Protein A domain D. In contrast, leucine at position 90 cannot form hydrogen bonds with N92 and R20, thereby removing the constraint on the side chain of R20. Consequently, the L90Q mutation may restore the interaction between R20 and Protein A.

To validate this hypothesis, we resolved the complex structures of VHH3-M1 and VHH3-M3 with Protein A domain D through co-crystallization, achieving resolutions of 1.49 Å and 2.0 Å, respectively. The interaction pattern between VHH3-M1 and Protein A closely resembles that of n501, with a root-mean-square deviation (RMSD) of 0.643 Å. This interaction is characterized by the formation of seven hydrogen bonds and two salt bridges (Supplementary Table 1). However, in VHH3-M1, the side chain of residue S17 is displaced, disrupting the hydrogen bond with A28 of Protein A domain D. Concurrently, the side chain of residue T77, also due to displacement, establishes a novel hydrogen bond with D39 of Protein A domain D. The structural alignment of VHH3-M3 with VHH3-M1 is highly conserved (0.164 Å), differing only at position 88, which is located outside the binding region and exerts minimal influence on binding affinity. Most notably, residues R20 and N92 not only engage in hydrogen bonding with the side chain of Q90 but also form hydrogen bonds with Protein A in both VHH3-M1 and VHH3-M3, corroborating our initial hypothesis.

### Redesigned sdAbs could be purified with Protein A resin

The objective of conferring Protein A binding to sdAbs is to facilitate label-free purification using commercial Protein A resin. To evaluate the efficiency of purifying these redesigned sdAbs with Protein A affinity chromatography, we divided the lysis supernatant into two equal aliquots and simultaneously purified them using both Ni-NTA and Protein A resins. Following elution with equal volumes, we compared the amounts of sdAb eluted (Fig. 5a). The results demonstrated that all mutants could be effectively captured and eluted by the Protein A resin (Fig. S4). Further analysis through grayscale scanning, using the Ni-NTA purification results as a reference, revealed that the purification efficiency of most mutants with Protein A resin exceeded 50%. Notably, at least one mutant from each sdAb exhibited purification efficiency comparable to that of Ni-NTA, thereby validating the reliability of the TFDesign-sdAb approach (Fig. 5b).

**Fig. 5.**
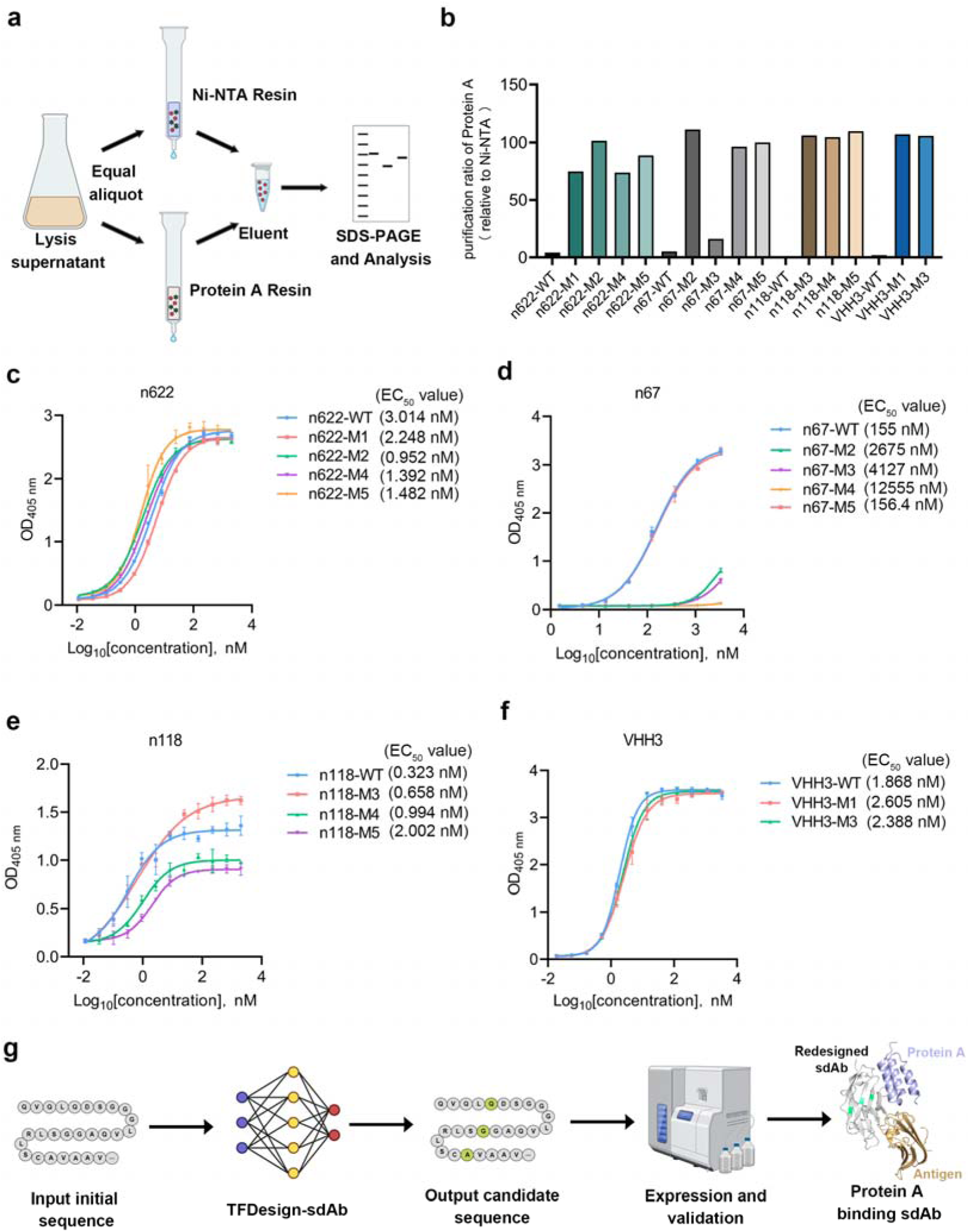
Efficient purification of optimized sdAbs using Protein A resin. **a.** Schematic diagram of sdAb purification using Ni-NTA and Protein A resins. The purified sdAbs were analyzed by SDS-PAGE **b.** Recovery rates of optimized sdAb mutants purified using Protein A resin, related to Ni-NTA purification. **c-f.** Four sdAb variants maintain high binding affinity to their cognate antigens. EC50 values are presented. The experiment was conducted with three replicates. **g.** Pipeline for sdAb design targeting Protein A with TFDesign-sdAb.

Additionally, we assessed the impact of the introduced mutations on the binding affinity of sdAbs to their cognate antigens. The results indicated that most mutants retained antigen-binding capabilities comparable to those of the wild type (Fig. 5c-f). This finding is consistent with the observation that the mutation sites are located within the framework region, which typically does not directly participate in antigen binding (Fig. S2).

In summary, TFDesign-sdAb enables the design of sdAbs that were previously incapable of binding Protein A to acquire this novel functionality, allowing for efficient label-free purification using Protein A affinity chromatography. This approach not only overcomes the limitations of traditional laboratory methods but also leverages artificial intelligence to achieve what was previously unfeasible—tailoring antibody binding properties through computational design. The pipeline simplifies the process by requiring only the input of the initial antibody sequence, followed by in silico design to generate candidate antibodies, predict their complex structures with antigens, and finally, perform expression verification to obtain the desired antibodies. This significantly reduces both the workload and costs compared to conventional experimental approaches (Fig. 5g).

## Discussion

Monoclonal antibodies (mAbs) have become essential therapeutic agents across various fields, including oncology, autoimmune disorders, and infectious diseases, owing to their high specificity and efficacy^27–29^. As the demand for next-generation antibodies with expanded applications continues to grow, engineering strategies must evolve to enhance their versatility while maintaining their core functions. The holy grail of antibody engineering lies in conferring novel functionalities without compromising native antigen-binding properties — a task demanding exquisite precision. Conventional approaches, such as phage display or site-directed mutagenesis, often yield suboptimal results due to the combinatorial complexity of antibody-protein interactions. For sdAbs, this challenge is magnified: their functional optimization requires modifying framework regions (FRs) rather than CDRs, as FRs dominate interactions with non-antigenic partners like Protein A.

In recent years, deep learning have been widely applied to protein design^12,30–35^, with antibody design emerging as one of the most challenging areas in this field^36,37^. Many efforts have been made to design the CDR regions of antibodies^38^ ^39^ ^40^ ^41^. However, these methods only optimize CDRs and do not alter the framework regions or their structural architecture. Although CDRs are crucial for binding to antigens, FRs also play an important role, especially in maintaining antibody stability and functionality. Especially in practical application scenarios, FRs support the function of antibodies. If FRs are inappropriate, antibodies may fail to function. Our approach, TFDesign-sdAb, addresses this gap by not only focusing on optimizing the CDRs but also taking the FRs into account. We collected antibody-antigen pairs from the SAbDab that interact with the antibody FR and trained a generative model for antibody FR design using a two-stage training process. Additionally, we employed the FR epitopes of Protein A from the complex structures as input to guide the optimization of specific regions of sdAbs. By considering both the CDRs and FRs in the design process, we can enhance the overall functionality of the antibody, enabling it to acquire new properties—such as the ability to bind to different targets—without sacrificing its original antigen-binding activity.

The downstream processing of therapeutic antibodies, particularly purification, represents a major cost in biopharmaceutical production. Among affinity-based purification methods, Protein A chromatography remains the gold standard due to its high specificity and scalability. However, the absence of an Fc domain in sdAbs precludes direct Protein A purification, necessitating affinity tags such as 6xHis, GST, or SUMO. While widely used, these tags can introduce immunogenicity, degradation issues, and additional processing steps that increase costs. Regulatory concerns surrounding tagged therapeutics further emphasize the need for streamlined, tag-free purification strategies to improve sdAb manufacturability. Therefore, we used Protein A as a case study to demonstrate how TFDesign-sdAb can confer sdAbs with the ability to bind a target they would not naturally recognize while preserving their original antigen specificity.

Existing strategies for enhancing Protein A binding in antibody fragments are limited, with one approach involving site-directed mutagenesis of nanobody or scFv constructs, which has achieved partial success in certain cases. However, the purification efficiency of these modified antibodies using Protein A remains significantly lower than that achieved with Ni-NTA resin. In our study, we redesigned a panel of distinct sdAbs, including fully human VHs and nanobodies, and selected the top five candidates from 1000 generated variants per sdAb. At least two candidates from each sdAb exhibited Protein A binding, achieving a 100% success rate, with at least one candidate per sdAb demonstrating purification efficiency comparable to Ni-NTA resin. These findings underscore the advantages of TFDesign-sdAb as a robust and generalizable approach for engineering sdAbs with new functional properties. Furthermore, TFDesign-sdAb can continuously optimize the sequence of sdAb without known structural information. This enables TFDesign-sdAb to be applied to a wider range of tasks and better meet actual needs.

The complex structures of antibodies with the domain D of Protein A were used to train the IgGM model, leading us to focus primarily on domain D in our subsequent research. We also explored the interactions between n501 and other domains of Protein A. Domain D shares approximately 80% sequence homology with the other four domains, and the 12 amino acid residues involved in binding are highly conserved. This conservation aligns with the similar EC50 values observed between n501 and different domains (Fig. S1), supporting the notion of similar interaction modes between sdAbs and other Protein A domains. We also identified that position 74 is crucial for sdAb binding to Protein A, with strict amino acid constraints. Mutations in surrounding residues can significantly influence binding. For example, in n67, variants M1 and M2 differ only a position 72, yet M1 shows minimal binding to Protein A. Similarly, in n118, a single mutation at S74 can enhance binding, but compared to our model-generated mutants, the affinity is reduced by 4.6-9.4 times (Fig. S6). This underscores the efficiency of our model compared to methods relying solely on mutations at key positions.

A key objective in antibody engineering is to retain the antibody’s ability to recognize its cognate antigen while enhancing other properties or introducing new functionalities. Although the CDRs of n501 do not interact with Protein A, we limited model-generated mutations to three to minimize potential impacts on antigen binding. Results indicated that most candidates with improved Protein A binding retained their ability to bind cognate antigens, with EC_50_ values for n622, n118, and VHH3 candidates comparable to those of the wild type. However, among n67 mutants with enhanced Protein A binding, only M5 maintained wild-type-like antigen affinity. This may be due to the simultaneous engagement of CDRs and framework regions in antigen binding, a possibility warranting further structural analysis. Overall, for each of the four antibodies, at least one candidate from our model enabled efficient purification using Protein A resin without compromising antigen binding.

In conclusion, TFDesign-sdAb provides a rapid, efficient, and cost-effective approach for designing sdAbs with tailored functionalities. By optimizing both CDRs and FRs, it enables the development of antibodies with novel binding properties while preserving their original antigen affinity. The successful use of this method to confer Protein A binding provides a proof of concept, highlighting the potential of AI-driven antibody design to broaden the functional capabilities of sdAbs for therapeutic and diagnostic purposes.

## Methods and materials

### Development of IgGM for sdAb Design

To develop IgGM model for the design of sdAb FRs, we employed a distinct training strategy consisting of two phases and utilized a different training set. In the first phase, we focused exclusively on structural prediction tasks, preserving the integrity of the original sequences and refraining from introducing noise. This phase concentrated on training the structural components using all antibody-antigen complex pairs from SAbDab^19^, which includes 8,355 antibody-antigen pairs clustered into 2,436 clusters using MMseqs2^42^ with 95% sequence identity.

In the second phase of training, we built upon the foundation established in the initial phase by focusing on sequence design. This phase integrated sequence recovery with structural prediction into a single model. Unlike the original IgGM model, the second phase utilized a subset of structural data from SAbDab, specifically including all antibody-antigen pairs where the antibody FR regions were in contact with the antigen (comprising 6,679 antibody-antigen pairs clustered into 2,030 clusters using MMseqs2 with 95% sequence identity). Furthermore, in the original IgGM training, the model was designed to predict CDRs, and noise was only added to the CDRs. For this task, however, we concentrated on optimizing the FR regions, focusing exclusively on those FR regions that were in contact with the antigen. For example, if the FR1 in the heavy chain contains amino acids that interact with the antigen, we would introduce noise to the sequence of FR1. This targeted approach allows for more effective optimization of the sdAb FRs in the context of antibody design.

As shown in Fig. S5a-b, we evaluated the structural prediction performance of the re-architected IgGM for the sdAb task (referred to as IgGM-FR) against the original IgGM (referred to as IgGM-CDR) using the n501-Protein A complex. The performance was assessed using DockQ for structural prediction and the Amino Acid Recovery rate (AAR) for paratope design. The DockQ score for IgGM-FR was 0.698, while the corresponding score for IgGM-CDR was only 0.032. At the interface between n501 and Protein A (distance < 8 Å), there are 19 amino acids involved; for the paratope recover, the recovery rate for IgGM-FR was 0.858, compared to just 0.078 for IgGM-CDR. Both sequence and structural metrics demonstrate the effectiveness of our retraining strategy, reflecting significant differences across various regions.

Additionally, we compared the structural prediction performance of IgGM with that of AlphaFold-3 for the VHH3-M1-Protein A and VHH3-M3-Protein A complexes, as illustrated in Fig. S5c-d. AlphaFold-3 incorrectly identified the paratope for VHH3-M1 and VHH3-M3 in relation to Protein A. In AlphaFold-3’s predictions, Protein A was incorrectly modeled to bind to CDR3 of VHH3, yielding DockQ scores of 0.03 and 0.018, respectively, which are significantly lower than IgGM’s predictions (0.816 for VHH3-M1 and 0.793 for VHH3-M3).

### Adaptation of A2Binder for sdAb ranking

Previous protein design methods^12,35^ typically utilize structure prediction tools (e.g., AlphaFold) as filters, employing metrics such as pTM, pLDDT, and PAE to screen candidate proteins. For antigen-antibody systems, some approaches have incorporated antigen-antibody docking^43^ and evaluated conformations to predict antigen-antibody binding affinity^44^ or changes in binding affinity^45,46^. However, when the accuracy of structure prediction methods for sdAb-Protein A complexes is low, these methods struggle to effectively filter candidates. Prior work has demonstrated that representations generated by protein language models can be utilized to predict protein functions, including thermostability and human protein-protein interactions (PPIs), as well as structures^47–51^. A2Binder has shown that representations generated by pretrained antibody language models (PALM) can be employed to predict the affinity of Fab antibodies for antigens^37^.

While sdAbs and mAbs share some sequence similarity, affinity data for sdAb-antigen interactions is considerably scarcer. To enhance the generalizability of our model, we trained it using both mAb and sdAb affinity data. To accommodate both types of inputs, we made simple modifications to the model’s data flow. Specifically, we duplicated the sdAb data and input it into the original VH and VL pre-trained models. This approach ensures that both sdAb and mAb data are processed through two pre-trained PALM models to extract sequence representations, maintaining a consistent distribution.

Additionally, A2Binder utilizes the BioMap dataset to train its model, which contains 1,706 antigenantibody (mAb) pairing data with affinity labels^52^. We further augmented this dataset with 86 antigen-sdAb and 653 antigen-mAb pairing affinity data from SAbDab ^19^, as well as 30 antigen-sdAb pairing affinity data measured in our laboratory, including n501-related data from alanine scanning. The ratio of sdAb to mAb data is 20:1, with sdAb data being unevenly distributed. To address this data imbalance, we reduced the frequency of mAb data in each epoch. Specifically, we clustered the mAb data based on the same antigen and selected only one mAb-antigen pair from a cluster for training in each epoch. This adjustment resulted in an effective training frequency ratio of sdAb to mAb data of approximately 4:1, significantly enhancing the model’s ability to generalize across different antibody types and improving its predictive performance for sdAb-antigen interactions.

### Expression and purification of human TNFα and CD16a

Recombinant human TFNα (hTNFα) was produced using Expi293 cells (ThermoFisher) via transient transfection with mammalian expression vector pcDNA3.1 (+), which contains the residues 77-233 (GenBank: AAA61198.1) and N-terminal signal peptide and 6xHis tag. Five days post-transfection, the supernatants were collected by centrifuging the culture at 3,900 rpm for 30 min and filtering through 0.45 *µ*m vacuum filter. Recombinant hTNFα was purified using Ni-NTA resin (Smart-Lifesciences) with elution buffer (10 mM Na2HPO4, 10 mM NaH2PO4 [pH 7.4], 500 mM NaCl, and 250 mM imidazole). The protein was then concentrated and buffer exchanged into PBS. Purity was analyzed by SDS-PAGE, and the protein concentration was determined using the NanoDrop 2000 spectrophotometer (ThermoFisher). Recombinant human CD16a (hCD16a) was also expressed in Expi293 cells (ThermoFisher) using the mammalian expression vector pSectag, which contains the residues 21-192 (GenBank: AAH33678.1) and N-terminal mouse Igκ signal peptide and C-terminal 6xHis tag. Additionally, two CD16A genes were linked with GPSMGSSSPS linker to form dimer. Five days post-transfection, hCD16a was purified using the same methods as described above.

### Expression and purification of sdAbs

The genes of n622, n118, n67, and VHH3, along with their mutants, were subcloned into the pComb3x vector, which includes an N-terminal OmpA signal peptide and a C-terminal 6xHis tag followed by a flag tag in series. Expression of these constructs was conducted in E. coli HB2151 at 30◦C for 14 hours, induced by 1 mM isopropyl β-D-1-thiogalactopyranoside (IPTG). The bacterial cells were harvested and then lysed with polymyxin B at 30°C for 45 minutes. After centrifugation at 8,000 rpm for 30 minutes, the supernatant was separated. For purification with Ni-NTA resin, the supernatant was applied to a Ni-NTA column following the manufacturer’s instructions (GE Healthcare). The resin was washed with buffer A (10 mM Na2HPO4, 10 mM NaH2PO4, pH 7.4, 500 mM NaCl, and 20 mM imidazole), and the proteins were eluted using buffer B (10 mM Na2HPO4, 10 mM NaH2PO4, pH 7.4, 500 mM NaCl, 250 mM imidazole). For purification with Protein A resin, the supernatant was loaded onto a Protein A resin column (Genscript) and washed with PBS. The bound proteins were eluted with 0.1 M glycine (pH 3.0) and neutralized by the addition of 1 mM Tris-HCl (pH 8.5). All harvests were buffer-exchanged to PBS, flash-frozen in liquid nitrogen, and stored at −80°C.

### Expression and purification of Protein A domains

The genes encoding the five domains of Protein A were constructed into the pET-28a(+) vector, each with a C-terminal 6×His tag (GenBank: WP_114967436.1; residues 37-92 for domain E, 93-153 for domain D, 154-211 for domain A, 212-269 for domain B, and 270-327 for domain C). The protein was expressed in the BL21 (DE3) strain and induced with 1 mM isopropyl-β-D-thiogalactoside for 16 hours at 30°C. The cell pellet was lysed in buffer A and purified as previously described. The finally purified protein was immediately used for crystallization. The remaining protein was flash-frozen in liquid nitrogen and stored at −80°C.

### Comparison of sdAb purification efficiency of different resins

To compare the purification efficiencies of Ni-NTA and Protein A resins for sdAb mutants, the following method was employed: sdAb was expressed, and the resulting supernatant was split into two aliquots. Each aliquot was then purified using either Ni-NTA or Protein A resin, as described previously. The eluates were concentrated and normalized to equal volumes before undergoing SDS-PAGE analysis. The gray-scale intensity of the sdAb bands was quantified using ImageJ software, with the Ni-NTA purified product serving as a control to determine the purification efficiency of the Protein A resin for sdAb mutants.

### Crystallization and data collection

Complexes of sdAb with Protein A domain D were formed by mixing each Protein at a 1:1 molar ratio overnight at 4°C, followed by isolation with SEC on a Superdex 75 10/300 GL column (GE Healthcare) in running buffer (20 mM HEPES, 100 mM NaCl, pH 7.5). All crystallization experiments were performed at 16◦C based on the sitting-drop method and were conducted using a Gryphon crystallization robot (Art Robbins Instruments, US) to add pool solution and protein samples. Specifically, 50 *µ*L of pool solution was dispensed into the designated well positions of a 96-well sitting-drop plate, while 0.3 *µ*L of protein sample and 0.3 *µ*L of pool solution were added to the corresponding drop positions on the same plate. After completing these operations, the 96-well sitting-drop plate was sealed with a membrane to ensure each well formed an independent. The complex of Protein A/n501 (17 mg/ml) with the molar ratio Protein A: nanobody 1:1 was used for crystal screening. Crystals of Protein A/n501 were grown at pH 5.5 in reservoir solution consisting of 25% w/v Polyethylene glycol 3350, 0.2M Magnesium chloride and 0.1M BIS-Tris. Cryoprotection was performed by adding glycerol to reservoir buffer at 20% concentration. The complex of Protein A/VHH3-M1 (12.3 mg/ml) with the molar ratio Protein A: nanobody 1:1 was used for crystal screening. Crystals of Protein A/VHH3-M1 were grown at pH 3.5 in reservoir solution consisting of 25% w/v Polyethylene glycol 3350 and 0.1M Citric acid. Cryoprotection was performed by adding glycerol to reservoir buffer at 20% concentration. The complex of Protein A/VHH3-M3 (9.7 mg/ml) with the molar ratio Protein A: nanobody 1:1 was used for crystal screening. Crystals of Protein A/VHH3-M3 were grown at 2.0 M Ammonium sulfate. Cryoprotection was performed by adding glycerol to reservoir buffer at 20% concentration.

The crystals were subsequently flash-frozen in liquid nitrogen prior to data acquisition. The X-ray diffraction data of the complex crystals were collected in beamline 10U2 at Shanghai Synchrotron Radiation Facility (SSRF).

### Determination and refinement of protein structure

Diffraction images were indexed and integrated with XDS^53^ or auto-processed by AutoPROC^54^ and Xia2_dials^55^ at Shanghai Synchrotron Radiation Facility (SSRF). The HKL files were obtained by molecular replacement using the Phaser program from CCP4 crystallography package^56^ with as the search model. Refmac^57^ and Phenix^58^ were used to refine the structure. The model was further adjusted by COOT^59^. The crystallographic parameters of Protein A/n501, Protein A/VHH3-M1, Protein A/VHH3-M3 complex are shown in supplementary Table 1. The related structural analysis figures were drawn by PyMOL ^60^.

### ELISA

The costar half-area high binding assay plates (Corning #3690) were coated with 100-200 ng/well of human TNFα, human CD16a, ZIKA E protein (40543-V08H, SinoBio) or human CEACAM5 (11077-H08H, SinoBio) in PBS overnight at 4◦C, followed by blocking with a PBS buffer containing 5% (w/v) bovine serum albumin (BSA) for 1 hour at 37◦C. SdAbs were then added in 3-fold serial dilutions. After incubation for 1.5 hours at 37◦C, the plate was washed three times with PBST (0.05% Tween 20 in PBS). Bound sdAbs were detected using an HRP-conjugated anti-flag monoclonal antibody (Sigma-Aldrich). Enzyme activity was measured by adding the substrate ABTS, and the signal was read at 405 nm using a Microplate Spectrophotometer (Biotek). The EC50 values were extrapolated by fitting the data to a sigmoidal dose-response curve with a variable slope using GraphPad Prism version 8.0.

For the detection of sdAb binding to Protein A and its five domains, the coating, blocking, and incubation were all performed using the same methods mentioned above. Additionally, a secondary antibody control group without the addition of sdAb was included. After the addition of ABTS, the plate was incubated at room temperature in the dark for 20 minutes to develop color, and the absorbance at 405 nm was read. The absorbance values of the experimental group were then adjusted by subtracting the values obtained from the secondary antibody control group.

### Biolayer interferometry (BLI) assay

The binding kinetics of sdAbs with Protein A were performed by BLI using Octet-Red96 device (Sartorius AG). Briefly, 3-fold serially diluted antibodies were incubated with ProA Biosensors (Sartorius AG). The assay mainly consisted of three steps: 1) baseline: 120 s with kinetics buffer (0.02% Tween 20 in PBS), 2) association: 300s with diluted antibodies, and 3) dissociation: 300 s with kinetics buffer. The sensors were then regenerated with pH 1.5 glycine for subsequent detection. A 1:1 binding model was used to calculate the exact KD with Data Analysis Software 11.1.

## List of Supplementary Materials

Fig. S1 to Fig. S6

Table S1 to Tables S2

## Acknowledgements

This work was supported by the National Natural Science Foundation of China (82394453, 92459301, 32270984), Science and Technology Commission of Shanghai Municipality (23XD1400800), Fund of Fudan University (24FCB09, FudanX24AI043), Shanghai Municipal Health Commission (GWVI-11.2-YQ46), and the Shanghai Frontiers Science Center of Pathogenic Microorganisms and Infection. We thank the staff from beamlines BL02U1 and BL18U1 at Shanghai Synchrotron Radiation Facility (SSRF) for assistance during data collection.

## Author contributions

T.Y., J.Y., Z.Y. and Y.W. initiated, planned and supervised the project. Y.K., J.S., F.W. and T.Z. performed most of the experiments with assistance from X.Z., Q.X., R.W., Y.S., Q.L., Y.W., X.G. and Y.Y. The manuscript was reviewed, commented and approved by all the authors.

## Conflict of Interest Statement

The authors declare that they have no conflict of interest.

## Notes

### Competing Interest Statement

The authors have declared no competing interest.

